# The Sphingosine-1-phosphate pathway is differentially activated in human gestational tissues

**DOI:** 10.1101/2025.06.02.657487

**Authors:** Magdaleena Naemi Mbadhi, Hideji Fujiwara, Ruth Gill, Kaci T. Mitchum, Cici Lin, Nandini Raghuraman, Antonina I. Frolova

**Author notes:** **Corresponding Authors:** 1. Antonina I. Frolova, Department of Obstetrics and Gynecology, Washington University in St. Louis, St. Louis, MO, USA; 2. Magdaleena Naemi Mbadhi, Department of Obstetrics and Gynecology, Washington University School of Medicine, St. Louis, MO, USA.

## Abstract

**BACKGROUND:** Dysregulated myometrial contractility contributes to obstetric complications. Sphingosine-1-phosphate (S1P) is an important inflammatory regulator in the myometrium and decidua, yet its metabolic dynamics during pregnancy are poorly characterized. This study aimed to profile the expression of S1P metabolic enzymes and receptors, and quantify sphingolipid metabolism in human gestational tissues across pregnancy.

**METHODS:** Myometrium, decidua parietalis, and chorioamnion were collected from women undergoing cesarean sections at term (≥37 weeks’ gestation) without labor (TNL), at term with labor (TL), and preterm (<37 weeks’ gestation) without labor (PTNL). Messenger RNA (mRNA) expression of S1P metabolic enzymes and receptors was assessed using quantitative polymerase chain reaction, while sphingolipids were quantified using targeted liquid chromatography-tandem mass spectrometry.

**RESULTS:** S1P metabolic enzymes and receptors were differentially expressed across gestational tissues. At TNL, *SPHK1* expression was significantly higher in the decidua parietalis than in the chorioamnion and myometrium. The myometrium exhibited the highest mRNA expression of S1P receptors (*S1PR1–4*) compared to the decidua and chorioamnion. At term, S1P was more abundant in the myometrium than in the decidua parietalis and chorioamnion. Both *SPHK1* and S1P were significantly increased in TL compared to TNL myometrium. S1P levels were higher in the myometrium at TNL compared to PTNL, while no significant differences were observed in the decidua and chorioamnion. Overall, sphingolipid metabolism was highest in the decidua and myometrium and lowest in the chorioamnion at term.

**CONCLUSION:** These findings reveal tissue-specific regulation of S1P metabolism and signaling in human gestational tissues, suggesting a therapeutic role of S1P in modulating myometrial contractility.

## Introduction

Parturition involves three key phenomena: the degradation of fetal membranes, synchronous myometrial contractions, and cervical ripening and dilation^1,2^. The physiological mechanisms governing human pregnancy and parturition are crucial for ensuring healthy pregnancy and labor outcomes. Preterm labor results from a premature activation of labor-inducing factors, triggering early myometrial contractility^3,4^. Conversely, prolonged gestation (delivery after 42 weeks) and labor arrest result from delayed onset or inadequate activation of labor-inducing factors, leading to insufficient myometrial contraction^5,6^. These labor abnormalities significantly increase the risk of neonatal morbidity and mortality and often necessitate delivery by cesarean section, which increases maternal morbidity rates^7–10^. To mitigate these labor complications and improve maternal and neonatal outcomes, we need to better understand the underlying mechanisms governing parturition.

The bioactive lipid mediator sphingosine-1-phosphate (S1P) was identified as a potential mediator of parturition, as it upregulated the expression of inflammatory markers associated with parturition in human myometrial cells^11^. S1P is generated by phosphorylation of sphingosine by the sphingosine kinase enzymes, SPHK1 and SPHK2^12^. SPHK1 is primarily located in the cytoplasmic compartment and translocates to the plasma membrane, where it is the principal generator of extracellular S1P^13^. In contrast, SPHK2 is located mainly in the cell nucleus, endoplasmic reticulum, and mitochondria and produces intracellular S1P that regulates gene transcription^14,15^. Once exported from the cell, S1P mediates its effects in an autocrine or paracrine fashion by binding and activating one of its five G protein-coupled receptors, S1P receptor 1-5 (S1PR1-5). S1P can then be recycled back to sphingosine by S1P phosphatases, S1PP1 and S1PP2, or irreversibly broken down by S1P lyase (S1PL) into non-sphingoid molecules^16^.

Accumulating evidence supports the involvement of S1P metabolic enzymes in pregnancy maintenance and parturition onset. For example, *Sphk1*-deficient mice exhibited increased decidual cell death, uterine hemorrhage, and decreased trophoblast cell invasion, resulting in impaired decidualization and pregnancy loss ^17^. SPHK1 activity and expression increase with advancing gestation in human decidua^18^, and its expression is higher in the myometrium during term labor than before labor onset^11^. Additionally, global inhibition of SPHK activity has been shown to inhibit lipopolysaccharide-induced preterm birth in mice^19,20^. Taken together, these studies suggest a potential physiological role for S1P in human parturition. However, the primary site of S1P production in the uterus has not been defined. We hypothesized that S1P synthesis is initiated in the decidua, triggering a signaling cascade that activates S1P-related enzymes and receptors in the myometrium to promote labor onset.

Here, we quantified the expression of enzymes involved in S1P synthesis and metabolism, as well as S1P receptors, in the fetal (chorioamniotic) membranes, decidua parietalis, and myometrium from women at term before or during labor. Furthermore, we used targeted mass spectrometry to quantify the concentrations of sphingolipid metabolites, including S1P, in these tissues. Finally, we compared the abundance of S1P and its two major precursors, sphinganine and sphingosine, in gestational tissues at preterm and at term pregnancy.

## Materials and Methods

### Study Population

This is a prospective cohort of nulliparous women who presented for delivery and underwent cesarean section at a single tertiary care institution between August 2022 and January 2024. Washington University in St. Louis Human Research Protection Office approved this study (IRB ID# 202205099). Multiple gestations and women with known HIV, hepatitis C or hepatitis B infections and gestational age ≥ 42 weeks were excluded.

Patients presenting for scheduled cesarean section at term (<u>≥</u>37 weeks gestation) were assigned to the term non-labor group (TNL, n=8), while those undergoing cesarean delivery in labor were assigned to the term labor group (TL, n=5). Patients undergoing unlabored cesarean section prior to 37 weeks gestation were assigned to the preterm non-labor group (PTNL, n=6). Patient clinical characteristics, including indications for cesarean delivery, are presented in **Table 1**.

**Table 1.** Patient characteristics.

| Characteristics | Term pregnancy |  | Preterm pregnancy |
| --- | --- | --- | --- |
|  | TNL (n=8) | TL (n=5) | PTNL (n=6) |
| Maternal age (yr) | 35.4 ± 5.1 | 31.4 ± 5.9 | 32.2 ± 7.0 |
| Gestational age (wk) | 38.6± 1.0 | 38.9 ± 1.4 | 33.9 ± 1.5 |
| Nulliparous | 3 (37.5) | 4 (80) | 0 |
| Length of labor (hrs) | N/A | 12.0 ± 7.3 | N/A |
| Indication for c/section: |  |  |  |
| Elective | 0 | 0 | 0 |
| Malpresentation | 3 (37.5) | 0 | 0 |
| Arrest of labor disorder | 0 | 3 (60) | 0 |
| Maternal indication/complications | 0 | 1 (20) | 0 |
| Non-reassuring fetal status | 0 | 0 | 0 |
| Repeat c/section | 4 (50.0) | 1 (20) | 5 (83.3) |
| Other | 1 (12.5) | 0 | 2 (33.3) |

Data represented are mean ± SD or n (%)

### Sample collection

Immediately after delivery of the neonate and placenta, an approximately 1.5 x 1.0 x 0.5 cm sample of myometrium was collected from the upper portion of the uterine incision site, placed into ice cold physiologic buffer solution in the operating room, and immediately transported to the lab. Once in the lab, the myometrial tissue was cut into ∼50 mg strips. The placenta was collected after delivery and transported to the lab. Decidua parietalis tissue was separated from the fetal membranes as previously described^21^. An approximately 50 mg piece of fetal membranes (chorioamnion) remaining following removal of the decidua parietalis was collected. All samples were flash frozen in liquid nitrogen within 30 minutes of collection and stored at −80 °C until further processing and analysis.

### RNA Isolation and Quantitative Polymerase Chain Reaction

Total RNA was isolated from the fetal membrane, decidua, and myometrium using the Direct-zol RNA Miniprep Kit (Zymo Research). RNA quality and quantity were assessed using the Nanodrop spectrophotometer. Total RNA (800 ng) was reverse-transcribed to cDNA using a high-capacity cDNA reverse transcription kit (Applied Biosystems) as per manufacturer’s protocol. cDNA was prepared, and mRNA gene expression was analyzed using real-time PCR (ABI7500; Life Technologies, MA, USA) using SYBR Green Master Mix as per manufacturer’s protocol. Human GAPDH was used as an internal control for gene expression normalization. Fold changes in gene expression were calculated using the 2^−ΔΔCT^ method. Primers used in this study were designed using PrimerQuest (Integrated DNA Technologies). Primer efficiencies were ≥85% and specificity was validated using the melt curve and electrophoresis analysis of the amplified product. Sequences of all primer sets used in the study are detailed in **Supplementary Table 1**.

### Sphingolipid analysis by liquid chromatography-tandem mass spectrometry (LC-MS/MS)

Four volumes of ice-cold water was added to each tissue (50 – 140mg) followed by homogenization in an Omni Bead Bluster (Perkin Elmer Inc) at 4°C for several minutes. The LC-MS/MS analysis was slightly modified from a previously published protocol^22^. Briefly, Sphingolipids (sphingoid bases, ceramides, and sphingomyelins (SM)) were extracted from 50 µL of the homogenate with 400 µL of (1:1) methanol/isopropyl alcohol containing internal standards (sphingosine-d7 10ng, sphinganine-d7, 5ng, S1P-d7 20ng, Cer(d18:1 17:0) 200ng, SM (d18: 1 17:0) 200 ng).

For sphingoid bases (sphingosine, sphinganine, and S1P) 10 µL of the extract was injected on a Water Xbridge C18 column (3.5 µm, 4.6 mm x 100 mm; Waters Corp.) connected to Shimadzu 20 ADX HPLC and API-4500 Qtrap tandem mass spectrometer (Applied Biosystem). The HPLC solvents were selected as A: 1% formic acid in water and B: 1% formic acid in 1:1 methanol/ACN. The solvent gradient was used from 70% B to 100% in 4 min at a flow rate of 1 mL/min and back to 70% in 2 min for the next injection. The positive ion multiple reaction monitoring (MRM) method was used to monitor sphingoid base analytes, including internal standards. Four-point calibration samples (0.8 ng/mL, 8 ng/mL, 20 ng/mL and 40 ng/mL) were prepared to obtain the absolute quantification data. All sphingoid base samples were injected twice to obtain the average data.

For ceramides and sphingomyelins (SM) ((Cer(d18:1 16:0), Cer(d18:1 18:0), Cer(d18:1 20:0), Cer(d18:1 22:0), Cer(d18:1 24:1), and Cer(d18:1 24:0) SM(d18:1 16:0), SM(d18:1 18:0), SM (d18:1 20:0), SM(d18:1 22:0), SM(d18:1 24:1), and SM(d18: 24:0)) 1 µL of the extract was injected on to an Agilent Zorbax Eclipse Plus C8 column (3.5 µm, 3.0 mm x 100 mm; Agilent) for HPLC/MS/MS analysis. The solvents were selected as A: 10 mM ammonium acetate in 30% ACN in water B: 10 mM ammonium acetate in 1:1 methanol/isopropyl alcohol. The solvent gradient was used from 65% B to 100% in 3 min at a flow rate of 1 mL/min and back to 65% in 2 min for the next injection. The positive ion MRM method was used for all ceramide and sphingomyelin (SM) detection, including the internal standards. 3- to 4-point calibration samples were also prepared for ceramide and SM absolute quantification. All samples were injected twice for the average data. All sphingolipids, including deuterated sphingolipids, were purchased from Avanti Polar Lipids Inc, 700 Industrial Park Drive. Alabaster, AL 35007. Analyst 5.1 software (Applied Biosystem) was used for this analysis.

### Statistical analysis

Statistical analyses were performed using GraphPad Prism software. The unpaired t-test with Welch correction assessed the statistical significance between two groups. For comparisons involving three groups, a one-way ANOVA was utilized, while a two-way ANOVA was applied for more than three groups, using Holm-Sidak’s method for multiple comparisons. Data was considered statistically significant at *p value ≤ 0.05, **p value ≤ 0.01, and ***p value ≤ 0.001.

## Results

### SPHK1 expression is highest in the decidua parietalis, and S1P receptor expression is highest in the myometrium at term non-labor

Previous works have analyzed the expression of S1P metabolic enzymes and receptors at term gestation in the decidua^18^ and myometrium^11^ independently. To further characterize S1P metabolism during parturition, we examined the expression of S1P synthesizing enzymes (*SPHK1* and *SPHK2*), degradation enzymes (*S1PP1* and *S1PL*), and receptors (S1PR1-4) in matched chorioamnion, decidua parietalis, and myometrium from patients at TNL. We found that mRNA for all enzymes and receptors analyzed was expressed in the tissues examined **(Figure 1)**. *SPHK1* mRNA was significantly more abundant in the decidua parietalis than in the chorioamnion and myometrium **(Figure 1A)**. *SPHK2* mRNA was highest in the myometrium **(Figure 1B)**. *S1PP1* mRNA was significantly more abundant in the decidua parietalis and myometrium than in the chorioamnion **(Figure 1C)**. S1PL mRNA levels were lowest in the chorioamnion, increased in the decidua parietalis, and showed the highest expression in the myometrium. **(Figure 1D)**.

**Fig 1.**
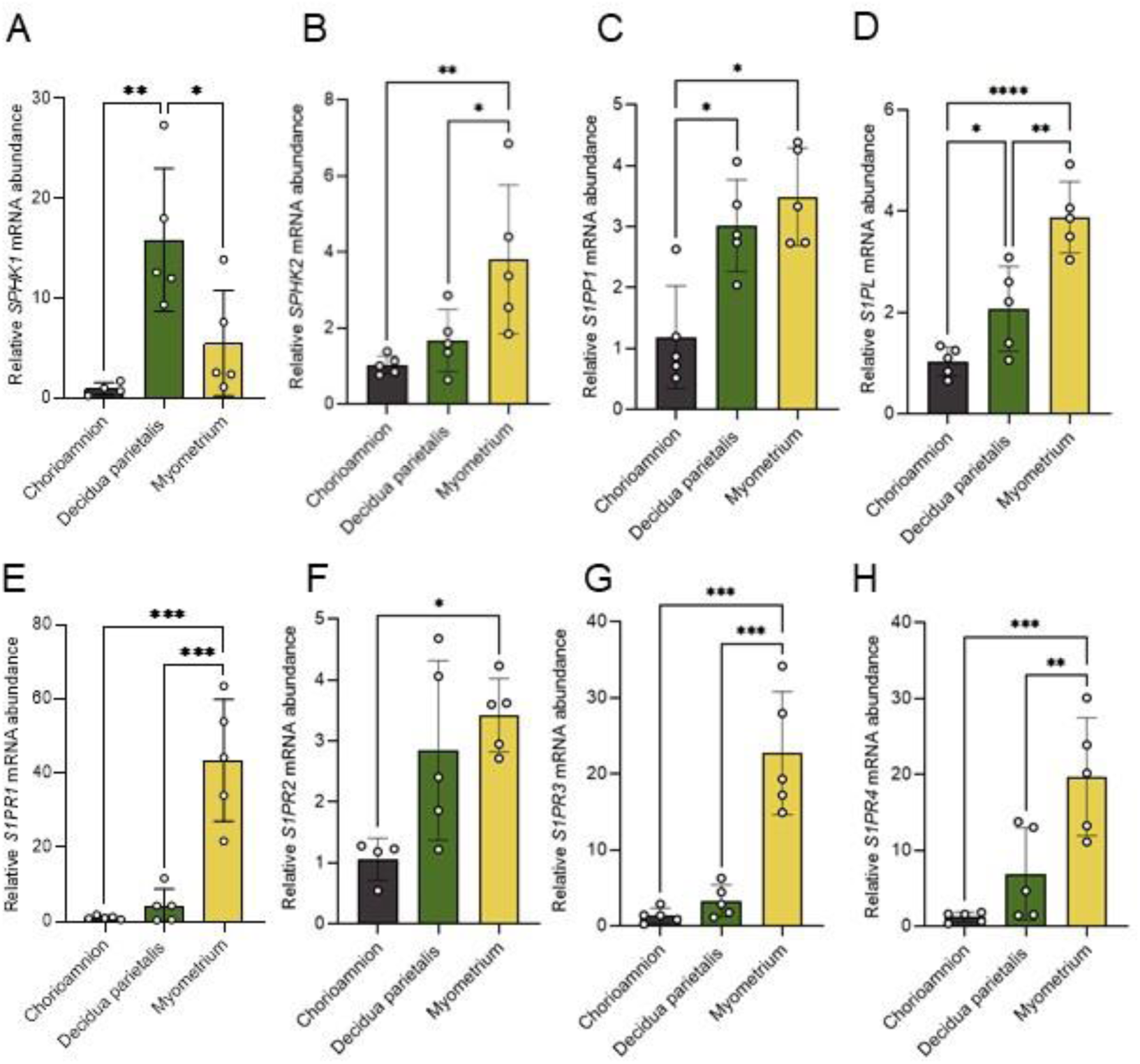
S1P metabolic enzyme and receptor mRNA expression in the human chorioamnion, decidua parietalis, and myometrium at term non-labor (TNL). mRNA expression of (A) sphingosine kinase 1 (SPHK1), (B) sphingosine kinase 2 (SPHK2), (C) sphingosine-1-phosphate phosphatase1 (S1PP1), (D) sphingosine-1-phosphate lyse (S1PL) in the human amnion, decidua parietalis, and myometrium during TNL were analyzed by quantitative real-time PCR (qPCR). Sphingosine-1-phosphate receptors (S1PRs), including (E) S1PR1, (F) S1PR2, (G) S1PR3, and (H) S1PR4 mRNA expressions, were also analyzed by qPCR in the human amnion, decidua parietalis, and myometrium at TNL. Presented are means ± SD of n=5. Statistical significance was determined by using one-way ANOVA and adjusted for multiple comparisons using the Holm-Sidak method (*P < 0.05; **P < 0.01; ***P < 0.001; ****P < 0.0001).

Expression of all four S1P receptors analyzed was highest in the myometrium and lowest in the chorioamnion **(Figure 1E-H)**. *S1PR1* mRNA was 40-fold higher in the myometrium than in the chorioamnion and 5-fold higher than in the decidua parietalis **(Figure 1E)**. *S1PR3* mRNA was 17-fold higher in the myometrium than in the chorioamnion and 6.8-fold higher than in the decidua parietalis **(Figure 1G)**.

### SPHK1 and S1PR3 are differentially expressed during labor in the human myometrium at term

Since S1P metabolic enzymes and receptors were most abundantly expressed in the decidua parietalis and myometrium, we compared their expression in these two tissues between patients at TNL and TL. Expression of IL8 was used as a marker of labor and notably increased in TL myometrial samples (**Supplementary Fig. 1**). In the decidua parietalis, expressions of *SPHK1, SPHK2, S1PP1, S1PL*, and *S1PR1-4* did not differ between TNL and TL **(Figure 2A, C)**. In the myometrium, *SPHK2, S1PP1, S1PL, S1PR1, S1PR2,* and *S1PR4* expression also did not differ between TNL and TL (**Figure 2B, D**). However, myometrial expression of *SPHK1* was 2.5-fold higher at TL than at TNL **(Figure 2B)**, whereas *S1PR3* expression was significantly higher at TNL than at TL **(Figure 2D)**.

**Fig 2.**
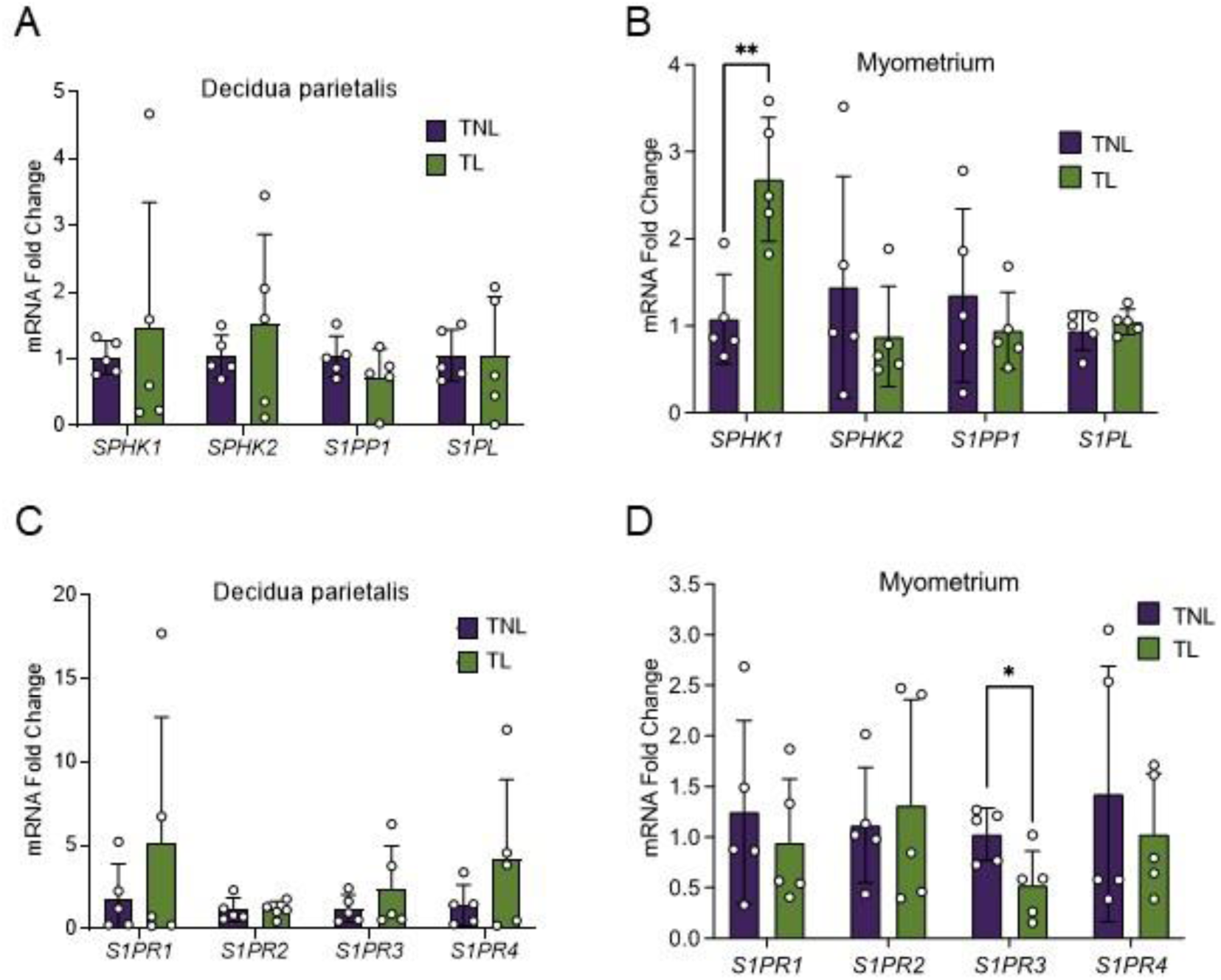
S1P metabolic enzyme and receptor mRNA expression in the human decidua parietalis and myometrium at term non-labor (TNL) and term-labor (TL). Expression of sphingosine kinase1 (SPHK1), sphingosine kinase2 (SPHK2), sphingosine-1-phosphate phosphatase1 (S1PP1), and sphingosine-1-phosphate lyse (S1PL) mRNA in the (A) decidua parietalis and (B) myometrium at TNL (n=5) and TL (n=5) were analyzed by quantitative real-time PCR (qPCR). Expression of sphingosine-1-phosphate receptors (S1PRs), including S1PR1, S1PR2, S1PR3, and S1PR4 mRNA in the (C) decidua parietalis and (D) myometrium at TNL and TL were analyzed by qPCR. Presented are means ± SD. Statistical significance was determined by using the unpaired t-test with Welch correction (*P < 0.05; **P < 0.01).

### Sphingolipid abundance within gestational tissues at term

To further explore the S1P synthesis pathway **(Figure 3A)** at term pregnancy, we used LC-MS/MS to quantify the abundance and distribution of 14 targeted sphingolipids in the chorioamnion, decidua parietalis, and myometrium at TNL and TL **(Figure 3B)**. Consistent with previous reports on sphingolipid abundance in the serum and plasma^23,24^, our results showed that sphingomyelins (SM) were also the most abundant sphingolipids in gestational tissues at term pregnancy. Specifically, SM C16:0 was the most abundant sphingomyelin at both TNL and TL.

**Figure 3.**
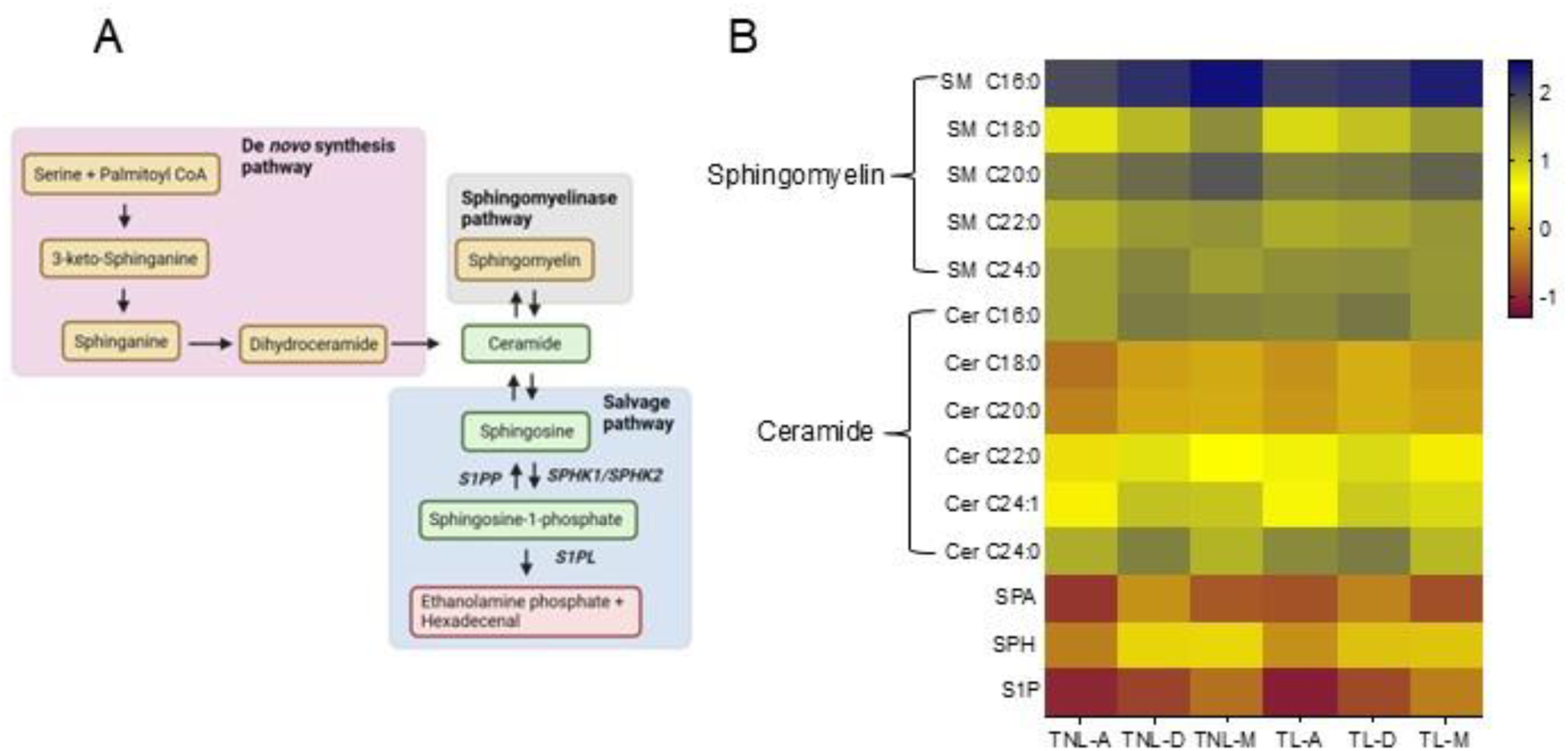
Distribution of sphingolipids in the chorioamnion, decidua parietalis, and myometrium at term non-labor (TNL) and term-labor (TL). (A) Schematic pathway of the sphingolipid synthesis. (B) Heatmap depicting the relative concentrations and distributions of 14 sphingolipids in the chorioamnion, decidua parietalis, and myometrium of TNL (n=8) and TL (n=5) quantified by mass spectrometry (LC-MS/MS). Sphingolipid relative abundance was log-transformed; the blue color bar represents high abundance, the yellow bar represents modest abundance, and the maroon color bar represents low abundance. A: chorioamnion; D: decidua parietalis; M: myometrium; TNL: term non-labor; TL: term labor; SM: sphingomyelin; Cer: ceramide; SPA: sphinganine; SPH: sphingosine; S1P: sphingosine-1-phosphate.

Ceramides, followed by sphingosine (SPH) - the direct S1P precursor - were the next most abundant sphingolipids in the chorioamnion and uterine tissues. Among ceramides, the fatty acid subclasses C16:0, C22:0, C24:0, and C24:1 were more abundant than C18:0 and C20:0 in all three tissues at both TNL and TL.

Sphinganine (SPA), the upstream precursor for ceramides and S1P in the *de novo* synthesis pathway, and S1P itself were the least abundant across all tissues. Specific concentrations of all sphingolipids across the different tissues at TNL and TL are provided in **Supplementary Tables 2-6**.

To identify the primary site of these bioactive lipid mediators, we compared SPA, ceramide, SPH, and S1P concentrations across the three gestational tissues and labor states. SPA concentrations were higher in the decidua parietalis than in the myometrium and chorioamnion **(Figure 4A)**. At TNL, ceramide-C16:0, -C18:0, -C20:0, -C22:0, and -C24:1 were higher in the decidua parietalis than in the chorioamnion, with no significant difference compared to the myometrium. At TL, ceramide-C18:0, -C20:0, and -C24:1 levels remained elevated in the decidua parietalis compared to the chorioamnion, and ceramide-C16:0, -C18:0, -C22:0, and -C24:1 were significantly more abundant in the decidua parietalis than in the myometrium (**Supplementary Tables 2 and 3)**.

**Figure 4.**
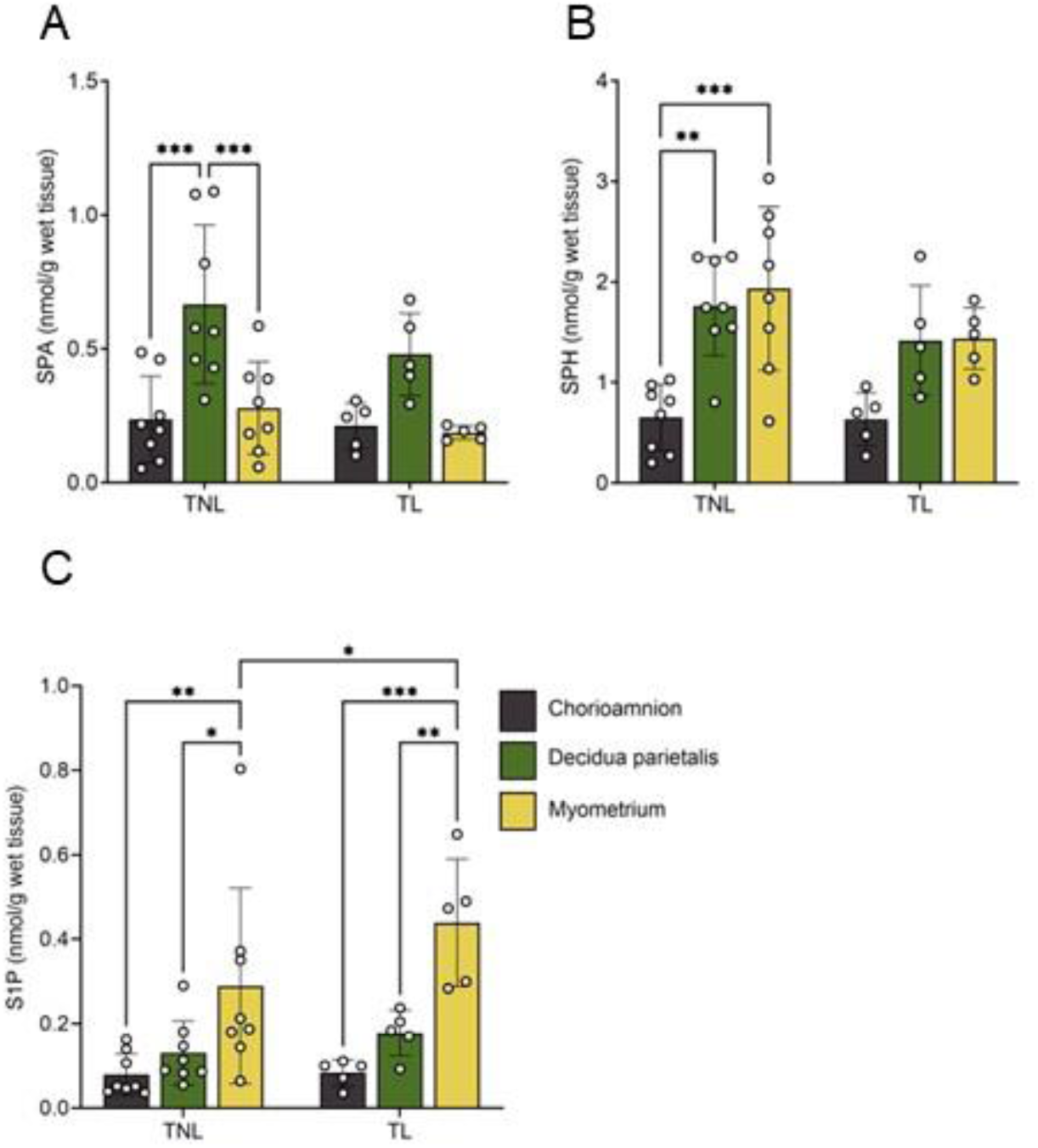
Bioactive sphingolipid concentrations in the human chorioamnion, decidua parietalis, and myometrium at term non-labor (TNL) and term labor (TL). (A) Sphinganine (SPA), (B) Sphingosine (SPH), and (C) sphingosine-1-phosphate (S1P) concentrations in the chorioamnion, decidua parietalis, and myometrium were analyzed by mass spectrometry (LC-MS/MS) in patients at TNL (n=8) and TL (n=5). Presented are means ± SD. Statistical significance was determined by using two-way ANOVA and adjusted for multiple comparisons using the Holm-Sidak method (*P < 0.05; **P < 0.01).

Similarly, SPH concentrations were comparable between the decidua parietalis and myometrium but lower in the chorioamnion at TNL, with no notable differences across tissues at TL **(Figure 4B)**. In contrast, S1P showed a distinct pattern, increasing from the chorioamnion to the decidua parietalis and reaching the highest levels in the myometrium at both TNL and TL **(Figure 4C)**. Additionally, S1P levels were higher in the myometrium at TL compared to TNL **(Figure 4C)**.

### S1P production is higher in the myometrium at term labor than at term non-labor

Given the differential abundance of S1P and its precursors (SPA and SPH) across gestational tissues, we next analyzed the ratios of S1P to its precursors as a proxy for turnover rate **(Figure 5A)**. The S1P:SPA ratio was highest in the myometrium compared to the chorioamnion and decidua parietalis at term **(Fig. 5B)**. Additionally, both the S1P:SPA and S1P:SPH ratios were significantly higher in the myometrium at TL than at TNL **(Fig. 5B and C)**, suggesting a higher S1P production rate in the myometrium during labor.

**Figure 5.**
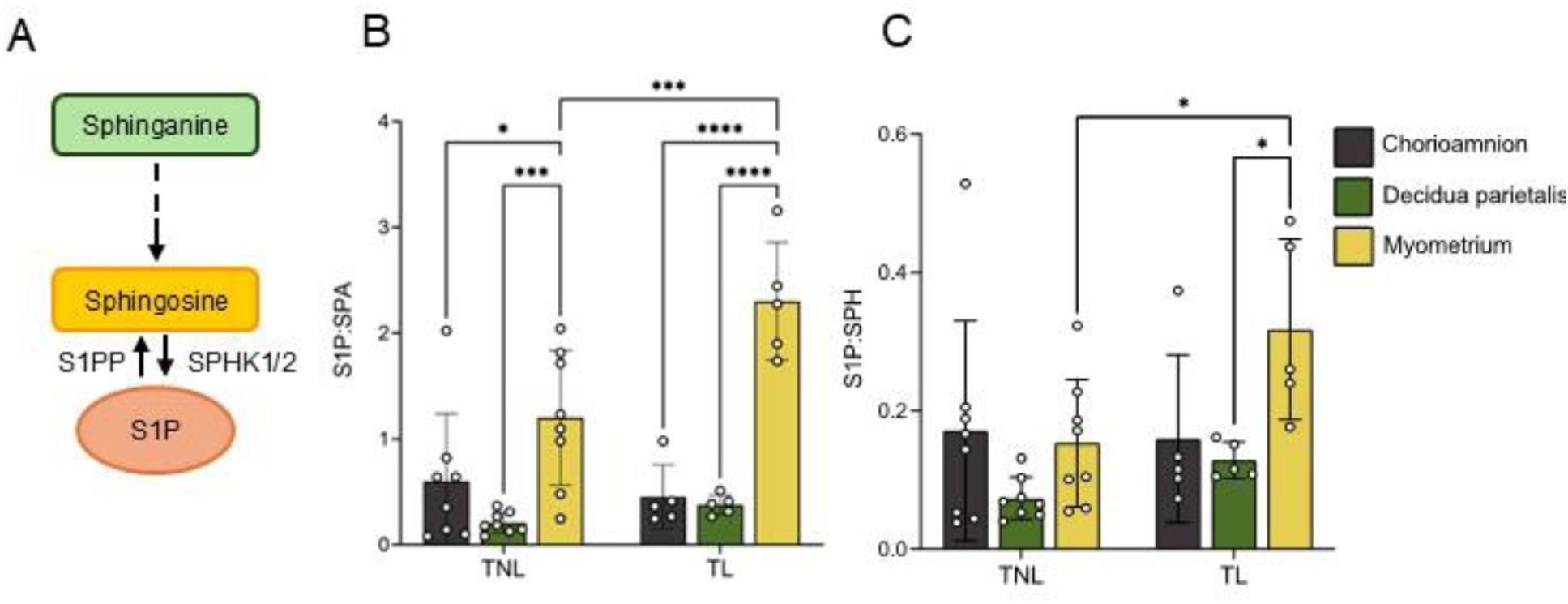
S1P to precursor ratios in the human chorioamnion, decidua parietalis, and myometrium at term non-labor (TNL) and term labor (TL). (A) Schematic pathway of S1P synthesis. (B) S1P-sphinganine (S1P:SPA) and (C) S1P-sphingosine (S1P:SPH) ratio in the human chorioamnion, decidua parietalis, and myometrium at TNL (n=8) and TL (n=5). Presented are means ± SD. Statistical significance was determined by using two-way ANOVA and adjusted for multiple comparisons using the Holm-Sidak method (*P < 0.05; **P < 0.01).

### Sphingolipid abundance in preterm gestational tissues

To explore whether similar sphingolipid metabolism patterns are present at preterm gestation, we analyzed SPA, SPH, and S1P levels in preterm non-labor (PTNL) tissues. Despite our analysis being limited to PTNL samples, SPA and SPH distribution mirrored those of TNL tissues. SPA was more abundant in the decidua parietalis than in the chorioamnion and myometrium (**Fig. 6A**), while SPH was similar between the decidua parietalis and myometrium and lowest in the chorioamnion **(Fig. 6B)**. In contrast to the distribution pattern at term, S1P abundance was similar between the decidua parietalis and myometrium and lowest in the chorioamnion **(Fig. 6C)**.

**Figure 6.**
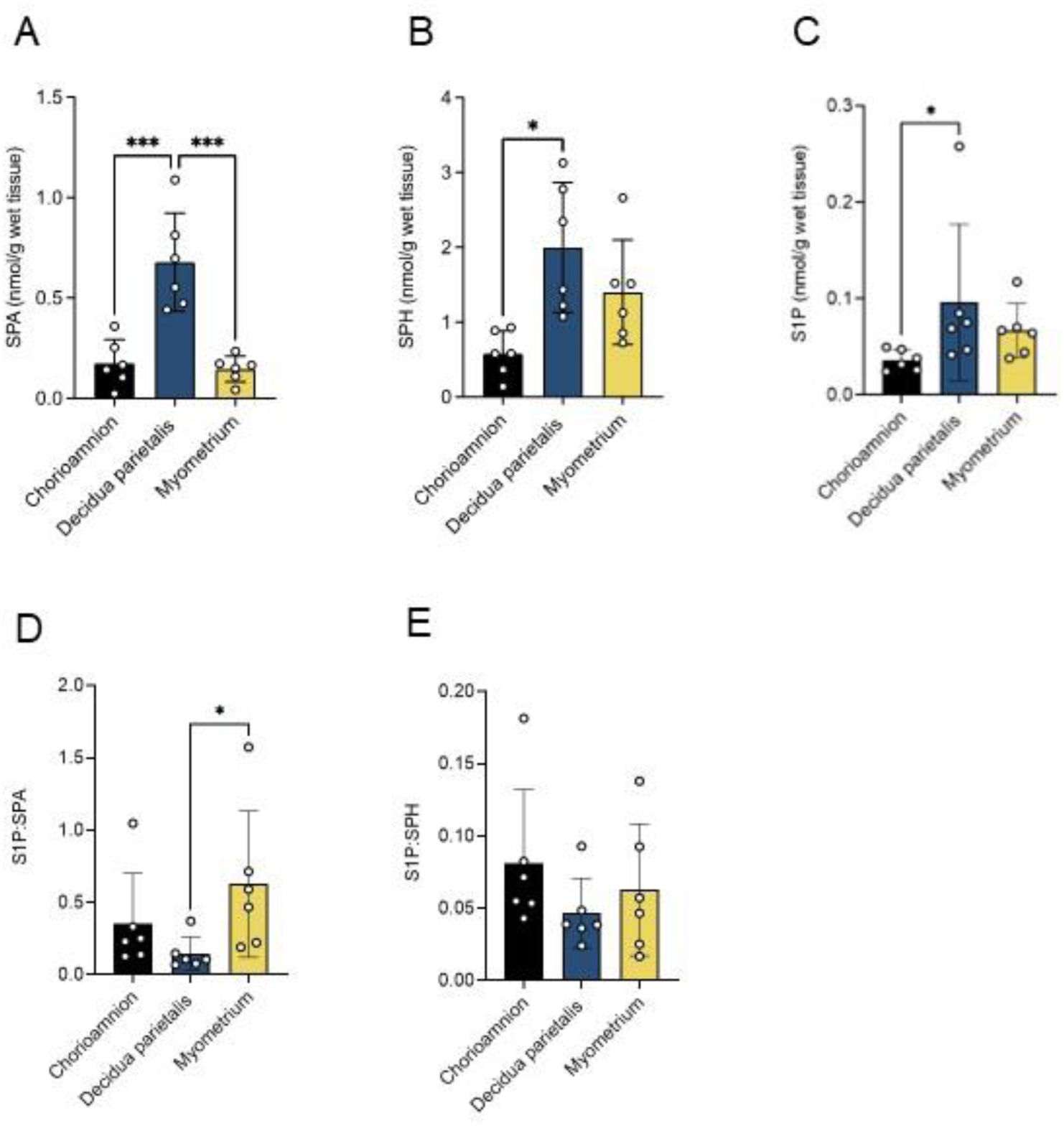
The bioactive sphingolipid concentrations in the human chorioamnion, decidua parietalis, and myometrium at preterm non-labor (PTNL). (A) Sphinganine (SPA), (B) Sphingosine (SPH), and (C) sphingosine-1-phosphate S1P concentrations in the chorioamnion, decidua parietalis, and myometrium were analyzed by mass spectrometry (LC-MS/MS) in patients at PTNL (n=7). (D) The S1P to sphinganine ratio (S1P:SPA) and (E) S1P to sphingosine ratio (S1P:SPH) in the chorioamnion, decidua parietalis, and myometrium was determined at PTNL (n=7). Presented are means ± SD. Statistical significance was determined using one-way ANOVA and adjusted for multiple comparisons using the Holm-Sidak method (*P < 0.05; **P < 0.01).

We next compared these metabolites between PTNL and TNL to evaluate gestational age-dependent differences. SPA and SPH abundances were similar between PTNL and TNL (**Supplementary Table 7)**. In contrast, S1P was significantly higher at TNL than at PTNL in the chorioamnion and myometrium (**Supplementary Table 7),** suggesting a gestational age-dependent increase in S1P.

Finally, we assessed S1P production and turnover by examining S1P:SPA and S1P:SPH ratios. At PTNL, the S1P:SPA ratio was higher in the myometrium than in the decidua parietalis **(Fig. 6D)**, suggesting increased S1P turnover activity via the *de novo* pathway. No significant differences in the S1P:SPH ratio were observed between tissues at PTNL (**Fig. 6E**). When comparing the gestational stages, the S1P:SPH ratio was higher in the myometrium at TNL than at PTNL (**Supplementary Table 7**), while the S1P:SPA ratio showed no difference between the two gestational stages.

## Discussion

This study investigated the expression of S1P metabolic enzymes, S1P receptors, and sphingolipid abundance in the human chorioamnion, decidua parietalis, and myometrium during preterm and term gestation. SPHK1, the rate-limiting enzyme for extracellular S1P production, exhibited the highest abundance in the decidua parietalis at TNL, and its expression in the myometrium increased with labor. Furthermore, S1P concentrations in the myometrium were significantly elevated at TL compared to TNL, indicating labor-dependent regulation of S1P in the tissue. Moreover, S1P levels were higher in the myometrium at TNL than at PTNL, suggesting that there is further gestational age-dependent regulation of S1P in the tissue. Finally, S1PR1-4 were most abundant in the myometrium, suggesting that this tissue is most responsive to S1P signaling.

Sphingolipid metabolites, including ceramide, sphingosine, and S1P, are known to be bioactive signaling molecules that regulate cell migration, differentiation, survival, and numerous other biological properties^25^. Although multiple sphingolipids have bioactive properties, S1P is the most potent and well-studied. Sphingolipid biosynthesis can occur via three distinct pathways, including the *de novo* pathway with sphinganine and ceramide as precursors, the sphingomyelinase pathway with sphingomyelins and ceramides as precursors, or the salvage pathway with sphingosine as the precursor^26,27^. These pathways all share a final rate-limiting step involving the phosphorylation of sphingosine by sphingosine kinases (SPHK1 and SPHK2) to generate S1P^26^. Thus, SPHK expression has been used as a surrogate of S1P production, concentration, and activity. Studies in sheep and rats have shown that SPHK1 expression increases in the uterus, including the myometrium, as gestation progresses^28,29^. Similarly, a human study has shown that SPHK1 protein and activity increase progressively in the decidua throughout pregnancy^18^.

With respect to parturition, SPHK1 expression was found to be higher in the human amniotic membrane at postpartum than at TNL^30^. Similarly, our group previously showed that SPHK1 gene expression was significantly higher in the myometrium of patients at TL compared to those at TNL^11^. Our current study supports and expands on these findings by quantifying the sphingolipid metabolites rather than solely relying on SPHK1 as a surrogate of S1P production. This approach is more comprehensive, as S1P levels are influenced by additional factors such as the presence and activity of S1P degradation enzymes and cellular transport mechanisms.

We showed that *SPHK1* gene expression increased in the myometrium at TL than at TNL, but this difference was not observed in the decidua. This spatial distinction suggests a specific association between S1P and the transition of the myometrium to a contractile state with the onset of labor, which is further supported by the higher abundance of S1P levels in the myometrium than in the decidua and chorioamnion at TL. These findings align with studies implicating S1P in preterm and term labor ^17,31–33^. One of the most compelling studies has shown that global inhibition of SPHK1/2, and therefore production of S1P, prevented inflammation-induced preterm labor in mice^19^. This was accompanied by a decrease in various proinflammatory cytokines, suggesting a significant role for S1P in regulating the inflammatory processes associated with labor onset. S1P has also been shown to directly induce the expression of cyclooxygenase-2 (COX-2) and other markers of labor in human and rat myometrial cells *in vitro*^11,28^.

We further found that S1P abundance was higher in the chorioamnion and myometrium at TNL than at PTNL. This finding is consistent with previous work that reported S1P was higher in the chorioamnion post-labor than at TNL^30^. The S1P:SPA ratio was not altered across gestational age, suggesting stable *de novo* production of S1P during pregnancy. However, the higher S1P:SPH ratio in the myometrium at TNL than at PTNL suggests increased S1P production due to increased SPHK1/2 activity with gestational age.

The primary source of S1P production within the gestational tissues remains a subject of ongoing inquiry. While some studies suggest that S1P produced in the amniotic tissue is a significant contributor during labor^30^, others have implicated the decidua^18^. Our sphingolipid metabolite profiling showed sphingomyelin, sphinganine, ceramide, and sphingosine levels were most abundant either in the decidua or equally abundant in the decidua and myometrium. We found consistently low levels of all sphingolipid metabolites, including *SPHK1*, in the chorioamnion compared to the maternal tissue. Instead, our data suggests that the decidua may play a substantial role in sphingolipid production, with elevated levels of sphingosine acting as a precursor for S1P. Our analysis, using matched samples, further demonstrated that *SPHK1* expression was higher in the decidua compared to the chorioamnion or myometrium at TNL, a crucial period when the decidua is priming for labor onset. Additionally, SM C16:0 was the most prominent sphingomyelin, while ceramide-C16:0 was the most prominent ceramide across all three gestational tissues at term pregnancy. Notably, ceramide-C16:0 was more abundant in the decidua than the chorioamnion at TNL, further illustrating the role of the decidua in regulating key sphingolipid metabolites involved in labor initiation. A notable exception was SM C16:0 and SM C18:0, which were found to be more abundant in the myometrium than in the decidua, particularly at TNL. While little is known about the function of sphingomyelin within tissues during pregnancy, decreased plasma levels of SM C16:0 and SM C18:0 during the first trimester have been associated with preeclampsia^23^. Research has also found increased plasma concentrations of ceramide-C16:0, a metabolite of SM C16:0, in patients experiencing early preterm labor compared to controls^34^. These findings suggest a potential role of these sphingolipid subclasses in gestational pathology, highlighting the necessity for further research on their functions in gestational tissue during labor.

S1P breakdown is regulated by the enzymes S1PP1 and S1PL. We found that the decidua and myometrium exhibited markedly higher levels of *S1PP1* than the chorioamnion at TNL. However, *S1PL* expression was more pronounced in the myometrium than in the decidua and chorioamnion at TNL. At TL, we did not observe any significant difference in either the S1P biosynthesis or degradation enzymes across the tissues, suggesting that most of the S1P enzymatic activity occurs before labor onset to accumulate the sufficient S1P levels necessary for signaling during parturition. The role of SPHK2 in pregnancy remains to be studied. However, we noted higher *SPHK2* expression in the myometrium than in the chorioamnion and decidua at TNL, likely due to transcriptional changes as the myometrium prepares for contraction.

Extracellular S1P exerts its biological effects by binding to one of its five G protein-coupled receptors, S1PR1-5, that initiate a diverse range of downstream signaling cascades^35^. We previously reported that S1P mediates its pro-inflammatory effects in human myometrial cells via S1PR3^11^, coinciding with findings that S1PR3 expression increased with gestation in human decidua^18^. Here, we found that *S1PR1-3* were more abundant in the myometrium than in the chorioamnion and decidua at term gestation, suggesting that the myometrium is the most responsive tissue to S1P signaling. There were no significant differences in *S1PR1-3* in the decidua or *S1PR1-2* expression in the myometrium between TL and TNL. An unexpected finding was decreased *S1PR3* expression in the myometrium at TL than TNL. The reduced receptor expression could be attributed to a negative feedback loop triggered by increased S1P signaling during the inflammatory process of labor, serving to maintain cellular homeostasis. It is crucial to note that S1P receptor signaling exhibits tissue- and cell-type-specificity, with the unique ability to elicit diverse responses depending on the environmental stimuli. Indeed, this is observed with S1PR1 expression, which increases only in chronic but not acute inflammation during inflammatory bowel disease^36^. Therefore, the complex interplay of labor and S1P signaling mechanisms may lead to differential regulation of the S1P receptors, including S1PR3.

While our study offers valuable insights, it is important to acknowledge several limitations. The sample size was relatively small, and we were unable to include preterm labor samples due to their limited availability. Additionally, our analyses were performed on whole-tissue samples rather than at the single-cell level, which may mask cell-specific differences in sphingolipid signaling. Future investigations utilizing single-cell approaches could provide a more detailed understanding of the distinct roles that sphingolipids play within individual cell types in gestational tissues. Despite these constraints, our study is strengthened by a comprehensive analysis of S1P metabolism and sphingolipid profiles in gestational tissues at three time points of gestation. Furthermore, we conducted our analyses using matched samples from the chorioamnion, decidua, and myometrium, which enhances the robustness and relevance of our findings.

Effectively managing labor complications, including preterm labor and labor dystocia, remains a challenge because current medications have off-target effects that result in adverse maternal and fetal outcomes. Our analysis revealed a striking tissue-specific disparity in S1P metabolites, enzymes, and receptors, with expression significantly higher in uterine tissues compared to fetal tissues (**Fig. 7**). This pronounced partitioning suggests that targeting S1P signaling could potentially bypass fetal tissue exposure, thereby minimizing fetal adverse effects associated with current systemic therapies.

**Figure 7.**
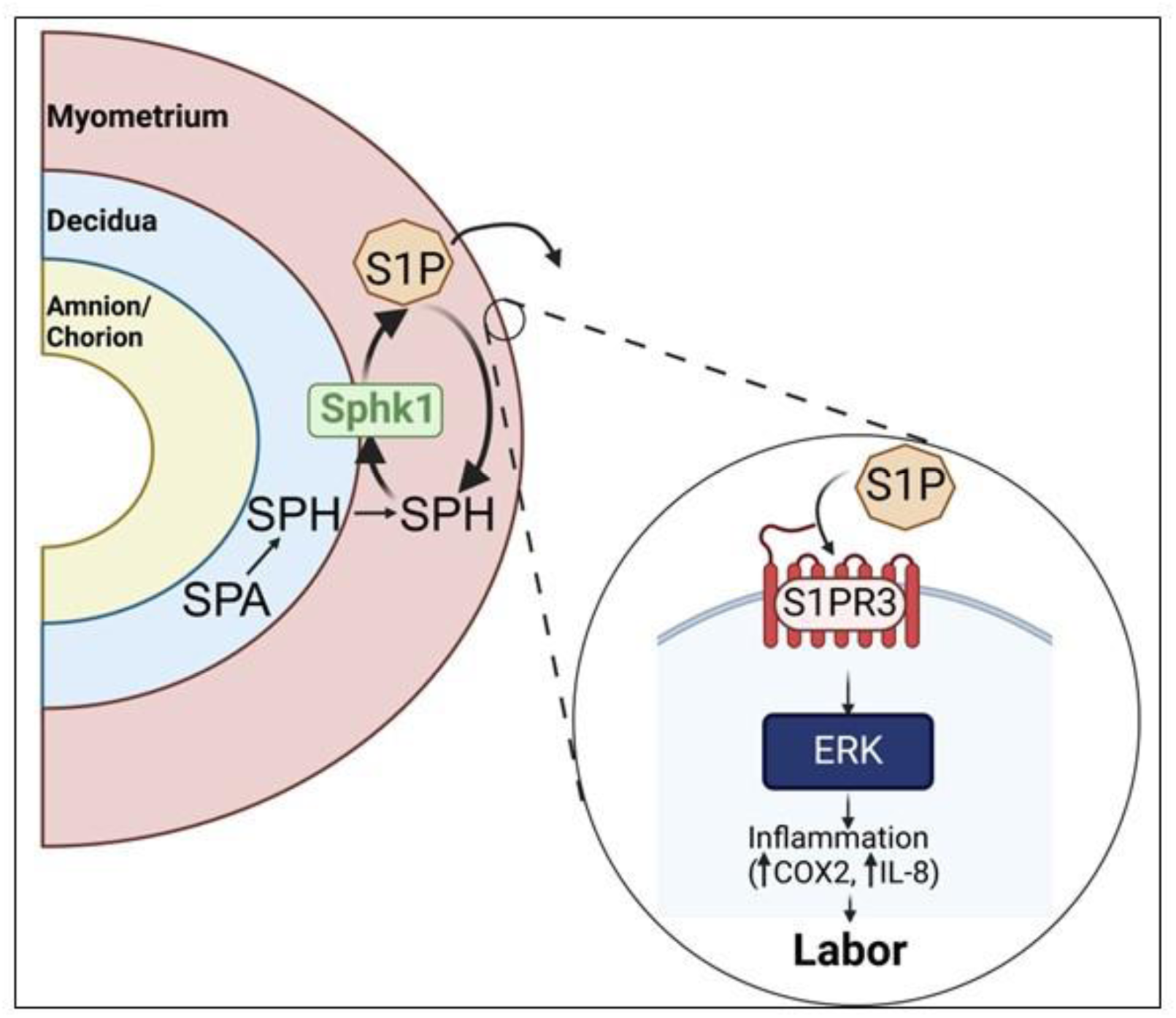
A proposed pathway for the metabolism and signaling of S1P in human gestational tissue. Sphingosine present in the decidua and myometrium leads to the production of S1P through the action of SPHK1. S1P predominantly exerts its effects on the myometrium by binding to S1PR3, subsequently triggering a downstream inflammatory cascade characterized by an elevation in COX2 and IL8 expression— processes associated with labor onset.

In conclusion, our study provides the first longitudinal, multi-tissue atlas of sphingolipid dynamics in the human gestational tissues. We demonstrate that S1P metabolism is differentially regulated across gestational tissues during pregnancy and labor. Notably, we showed that S1P production and signaling are highly modulated in the myometrium during labor. This tissue-specific expression pattern highlights S1P signaling as a precision therapeutic target for labor-related complications, with potentially reduced off-target fetal side effects. This work lays the foundation for future studies to investigate the potential efficacy and safety of S1P-based therapies in gestational models for modulating the myometrium during labor.

## Supporting information

Specific concentrations of all sphingolipids across the different tissues at TNL and TL are provided in Supplementary Tables 2-6.

Expression of IL8 was used as a marker of labor and notably increased in TL myometrial samples (Supplementary Fig. 1)

## Acknowledgement

We thank Dr Deborah Frank for the invaluable editorial assistance. We thank the Washington University School of Medicine ReProBANK for tissue collection and processing.

## Authorship contribution

M.N.M and A.I.F conceptualization, methodology, and formal analysis; A.I.F funding acquisition; M.N.M, H.F, R.G, K.T.M, and C.L investigation; A.I.F supervision; M.N.M visualization; M.N.M writing – original draft; M.N.M, N.R, and A.I.F writing – review and editing

## Supp

**Supplemental Fig 1. An inflammatory marker of parturition in the myometrium of term non-labor (TNL) and term-labor (TL) human tissue.** mRNA expression of IL-8 in the myometrium of TNL (n=5) and TL (n=5) patients was analyzed by quantitative real-time PCR (RT-PCR). Presented are means ± SD. Statistical significance was determined by using multiple unpaired t-test with Welch correction (*P < 0.05).

