## Supplementary material for "The Sphingosine-1-phosphate pathway is differentially activated in human gestational tissues": Specific concentrations of all sphingolipids across the different tissues at TNL and TL are provided in Supplementary Tables 2-6.

**Supplemental Table 1.** Primer sequences for real-time PCR

| Gene | Forward primer | Reverse primer |
|---|---|---|
| SPHK1 | 5’-GGCAGGCATATGGAGTATGAA-3’ | 5’-CCACTGCAAACACACCTTTC-3’ |
| SPHK2 | 5’-CCCGGTTGCTTCTATTGGT-3’ | 5’-CCTTCAACCTCATCCAGACAG-3’ |
| S1PR1 | 5’-CACTCTGACCAACAAGGAGATG-3’ | 5’-GATGATGGGTCGCTTGAATTTG-3’ |
| S1PR2 | 5’-ATCGTGCTAGGCGTCTTTATC-3’ | 5’-CCTCTACAAAGCCCACTACTTT-3’ |
| S1PR3 | 5’-CTCTACGCACGCATCTACTTC-3’ | 5’-AACACGCTCACCACAATCA-3’ |
| S1PR4 | 5’-CGTCTTTGGCTCCAACCT-3’ | 5’-CTGCGGAAGGAGTAGATGATG-3’ |
| S1PP1 | 5' -GCCGCTGGCAGTACCCTCT-3' | 5' -GTTGAAGTTGTCAATCAGGTCCACA-3' |
| S1PL | 5'- TCCGGCGTGAGGAGAGTCTG-3' | 5'- GCTGCCAGGGCTCATACTTG-3' |
| IL-8 | 5’-GCCTTCCTGATTTCTGCAGC-3’ | 5’-CGCAGTGTGGTCCACTCTCA-3’ |
| COX2 | 5’-TACTGGAAGCCAAGCACTTT-3’ | 5’-GGACAGCCCTTCACGTTATT-3’ |
| GAPDH | 5’-GGTGTGAACCATGAGAAGTATGA-3’ | 5’-GAGTCCTTCCACGATACCAAAG-3’ |

**Supplemental Table 2.** Concentrations of sphingolipids in the chorioamnion and uterine tissue of TNL patients

| **Sphingolipids (nmol/g tissue)** | **Tissue type** | | | **Adjusted p-Value** | | |
|---|---|---|---|---|---|---|
| | **Chorioamnion (n=8)** | **Decidua (n=8)** | **Myometrium (n=8)** | **AvD** | **AvM** | **DvM** |
| SM C16:0 | 91.9 ± 28.6 | 144 ± 14.6 | 239 ± 20.4 | 0.0003 *** | <0.0001**** | <0.0001**** |

| | | | | | | |
|---|---|---|---|---|---|---|
| SM C18:0 | 6.32 ± 1.60 | 13.5 ± 2.33 | 28.7 ± 5.42 | 0.0007*** | <0.0001 **** | <0.0001 **** |
| SM C20:0 | 34.1 ± 12.5 | 51.9 ± 14.8 | 72.0 ± 11.2 | 0.0126* | <0.0001 **** | 0.0119* |
| SM C22:0 | 14.4 ± 2.77 | 23.0 ± 5.21 | 26.2 ± 5.07 | 0.0061** | 0.0006*** | 0.2188 |
| SM C24:0 | 19.9 ± 4.63 | 33.8 ± 9.01 | 21.8 ± 5.84 | 0.0024** | 0.5822 | 0.0052** |
| Cer C16:0 | 20.5 ± 7.09 | 41.0 ± 19.0 | 37.0 ± 16.8 | 0.0252* | 0.0558 | 0.5639 |
| Cer C18:0 | 0.34 ± 0.06 | 0.79 ± 0.32 | 0.92 ± 0.28 | 0.0027** | 0.0004*** | 0.2760 |
| Cer C20:0 | 0.46 ± 0.10 | 0.92 ± 0.29 | 1.00 ± 0.39 | 0.0030** | 0.001** | 0.4964 |
| Cer C22:0 | 2.48 ± 1.03 | 7.04 ± 3.06 | 4.05 ± 1.86 | 0.0027** | 0.1817 | 0.0339* |
| Cer C24:0 | 17.3 ± 6.78 | 37.5 ± 16.2 | 15.4 ± 5.85 | 0.0275* | 0.8038 | 0.0244* |
| Cer C24:1 | 3.39 ± 1.52 | 12.2 ± 4.64 | 11.1 ± 3.27 | 0.0002*** | 0.0006*** | 0.5177 |
| SPA | 0.24 ± 0.16 | 0.67 ± 0.30 | 0.28 ± 0.17 | 0.0003*** | 0.6443 | 0.0006*** |
| SPH | 0.65 ± 0.33 | 1.76 ± 0.49 | 1.94 ± 0.81 | 0.0011** | 0.0004*** | 0.5293 |

| | | | | | | |
|---|---|---|---|---|---|---|
| S1P | 0.08 ± 0.05 | 0.13 ± 0.08 | 0.29 ± 0.23 | 0.3922 | 0.0051** | 0.0259* |

**Supplementary Table 2.** Targeted sphingolipid abundance in the chorioamnion, decidua parietalis, and myometrium at term non-labor. Statistical significance was determined by two-way ANOVA, and significant differences were followed up with Holm-Šídák's multiple comparisons test. Data represented are mean ± SD. A: chorioamnion; D: decidua parietalis; M: myometrium; TNL: term non-labor; SM: sphingomyelin; Cer: ceramide; SPA: sphinganine; SPH: sphingosine; S1P: sphingosine-1-phosphate.

**Supplemental Table 3**. Concentrations of sphingolipids in the amnion and uterine tissue of TL patients

| **Sphingolipids (nmol/g tissue)** | **Tissue type** | | | **Adjusted p-Value** | | |
|---|---|---|---|---|---|---|
| | **Chorioamnion (n=5)** | **Decidua (n=5)** | **Myometrium (n=5)** | **AvD** | **AvM** | **DvM** |
| SM C16:0 | 109 ± 19.3 | 129 ± 25.6 | 191 ± 29.4 | 0.1065 | <0.0001 **** | 0.0001*** |
| SM C18:0 | 7.67 ± 1.45 | 11.7 ± 2.64 | 22.7 ± 5.70 | 0.0300* | <0.0001 **** | <0.0001 **** |
| SM C20:0 | 38.9 ± 13.1 | 45.0 ± 17.1 | 60.8 ± 20.8 | 0.3502 | 0.0096** | 0.0469* |
| SM C22:0 | 16.8 ± 3.97 | 19.6 ± 5.80 | 24.2 ± 3.89 | 0.2735 | 0.0246* | 0.1510 |
| SM C24:0 | 27.4 ± 3.92 | 29.1 ± 7.58 | 24.0 ± 4.68 | 0.6161 | 0.5471 | 0.3802 |

| | | | | | | |
|---|---|---|---|---|---|---|
| Cer C16:0 | 31.8 ± 9.28 | 43.6 ± 15.7 | 24.3 ± 4.85 | 0.1970 | 0.2888 | 0.0364* |
| Cer C18:0 | 0.60 ± 0.09 | 1.01 ± 0.34 | 0.72 ± 0.15 | 0.0072** | 0.3144 | 0.0415* |
| Cer C20:0 | 0.66 ± 0.19 | 1.01 ± 0.27 | 0.79 ± 0.14 | 0.0299* | 0.2978 | 0.1611 |
| Cer C22:0 | 5.28 ± 2.58 | 7.89 ± 2.65 | 2.91 ± 0.74 | 0.0657 | 0.0657 | 0.0013** |
| Cer C24:0 | 30.3 ± 8.02 | 42.1 ± 19.7 | 13.9 ± 4.23 | 0.1245 | 0.0780 | 0.0042 ** |
| Cer C24:1 | 4.65 ± 0.95 | 10.7 ± 4.32 | 7.68 ± 1.53 | 0.0068** | 0.1650 | 0.1650 |
| SPA | 0.21 ± 0.09 | 0.48 ± 0.15 | 0.19 ± 0.03 | 0.0578 | 0.8274 | 0.0536 |
| SPH | 0.63 ± 0.27 | 1.42 ± 0.55 | 1.44 ± 0.31 | 0.0885 | 0.0885 | 0.9606 |
| S1P | 0.08 ± 0.03 | 0.18 ± 0.05 | 0.44 ± 0.15 | 0.2199 | 0.0003*** | 0.0039** |

**Supplementary Table 3.** Targeted sphingolipid abundance in the chorioamnion, decidua parietalis, and myometrium at term labor. Statistical significance was determined by two-way ANOVA, and significant differences were followed up with Holm-Šídák's multiple comparisons test. Data represented are mean ± SD. A: chorioamnion; D: decidua parietalis; M: myometrium; TL: term labor; SM: sphingomyelin; Cer: ceramide; SPA: sphinganine; SPH: sphingosine; S1P: sphingosine-1-phosphate.

**Supplemental Table 4**. Concentrations of sphingolipids in the chorioamnion of TNL vs TL patients

| Sphingolipids (nmol/g tissue) | Labor type | | |
|---|---|---|---|
| | **TNL (n=8)** | **TL (n=5)** | **Adjusted p-Value** |
| SM C16:0 | 91.9 ± 28.6 | 109.2 ± 19.3 | 0.2594 |
| SM C18:0 | 6.32 ± 1.60 | 7.67 ± 1.45 | 0.5626 |
| SM C20:0 | 34.1 ± 12.5 | 38.9 ± 13.1 | 0.6277 |
| SM C22:0 | 14.4 ± 2.77 | 16.8 ± 3.97 | 0.4248 |
| SM C24:0 | 19.9 ± 4.63 | 27.4 ± 3.92 | 0.0691 |
| Cer C16:0 | 20.5 ± 7.09 | 31.8 ± 9.28 | 0.1913 |
| Cer C18:0 | 0.34 ± 0.06 | 0.60 ± 0.09 | 0.0997 |
| Cer C20:0 | 0.46 ± 0.10 | 0.66 ± 0.19 | 0.2062 |
| Cer C22:0 | 2.48 ± 1.03 | 5.28 ± 2.58 | 0.0522 |
| Cer C24:0 | 17.3 ± 6.78 | 30.3 ± 8.02 | 0.0902 |
| Cer C24:1 | 3.39 ± 1.52 | 4.65 ± 0.95 | 0.5229 |
| SPA | 0.24 ± 0.16 | 0.21 ± 0.09 | 0.8201 |
| SPH | 0.65 ± 0.33 | 0.63 ± 0.27 | 0.9491 |
| S1P | 0.08 ± 0.05 | 0.08 ± 0.03 | 0.9508 |

**Supplementary Table 4.** Targeted sphingolipid abundance in the chorioamnion at term non-labor and term labor. Statistical significance was determined by two-way ANOVA, and significant differences were followed up with Holm-Šídák's multiple comparisons test. Data represented are mean ± SD. TNL: term non-labor; TL: term labor; SM: sphingomyelin; Cer: ceramide; SPA: sphinganine; SPH: sphingosine; S1P: sphingosine-1-phosphate.

**Supplemental Table 5.** Concentrations of sphingolipids in the decidua parietalis of TNL vs TL patients

| **Sphingolipids (nmol/g tissue)** | **Labor type** | | |
|---|---|---|---|
| | **TNL (n=8)** | **TL (n=5)** | **Adjusted p-Value** |
| SM C16:0 | 144 ± 14.6 | 129 ± 25.6 | 0.3180 |
| SM d18:1 18:0 | 13.5 ± 2.33 | 11.7 ± 2.64 | 0.4517 |
| SM C20:0 | 51.9 ± 14.8 | 45.0 ± 17.1 | 0.4785 |
| SM C22:0 | 23.0 ± 5.21 | 19.6 ± 5.80 | 0.2423 |
| SM C24:0 | 33.8 ± 9.01 | 29.1 ± 7.58 | 0.2408 |
| Cer C16:0 | 41.0 ± 19.0 | 43.6 ± 15.7 | 0.7651 |
| Cer C18:0 | 0.79 ± 0.32 | 1.01 ± 0.34 | 0.1453 |
| Cer C20:0 | 0.92 ± 0.29 | 1.01 ± 0.27 | 0.5465 |
| Cer C22:0 | 7.04 ± 3.06 | 7.89 ± 2.65 | 0.5395 |
| Cer C24:0 | 37.5 ± 16.2 | 42.1 ± 19.7 | 0.5332 |
| Cer C24:1 | 12.2 ± 4.64 | 10.7 ± 4.32 | 0.4453 |
| SPA | 0.67 ± 0.30 | 0.48 ± 0.15 | 0.0863 |
| SPH | 1.76 ± 0.49 | 1.42 ± 0.55 | 0.2601 |
| S1P | 0.13 ± 0.08 | 0.18 ± 0.05 | 0.5227 |

**Supplementary Table 5.** Targeted sphingolipid abundance in the decidua parietalis at term non-labor and term labor. Statistical significance was determined by two-way ANOVA, and significant differences were followed up with Holm-Šídák's multiple comparisons test. Data represented are mean ± SD. TNL: term non-labor; TL: term labor; SM: sphingomyelin; Cer: ceramide; SPA: sphinganine; SPH: sphingosine; S1P: sphingosine-1-phosphate.

**Supplemental Table 6**. Concentrations of sphingolipids in the myometrium of TNL vs TL patients

| Sphingolipids (nmol/g tissue) | Labor type | | |
|---|---|---|---|
| | **TNL (n=8)** | **TL (n=5)** | **Adjusted p-Value** |
| SM C16:0 | 239 ± 20.4 | 191 ± 29.4 | 0.0039** |
| SM d18:1 18:0 | 28.7 ± 5.42 | 22.7 ± 5.70 | 0.0145** |
| SM C20:0 | 72.0 ± 11.2 | 60.8 ± 20.8 | 0.2586 |
| SM C22:0 | 26.2 ± 5.07 | 24.2 ± 3.89 | 0.4987 |
| SM C24:0 | 21.8 ± 5.84 | 24.0 ± 4.68 | 0.5871 |
| Cer C16:0 | 37.0 ± 16.8 | 24.3 ± 4.85 | 0.1412 |
| Cer C18:0 | 0.92 ± 0.28 | 0.72 ± 0.15 | 0.1901 |
| Cer C20:0 | 1.00 ± 0.39 | 0.79 ± 0.14 | 0.2003 |
| Cer C22:0 | 4.05 ± 1.86 | 2.91 ± 0.74 | 0.4153 |
| Cer C24:0 | 15.4 ± 5.85 | 13.9 ± 4.23 | 0.8396 |
| Cer C24:1 | 11.1 ± 3.27 | 7.68 ± 1.53 | 0.0938 |
| SPA | 0.28 ± 0.17 | 0.19 ± 0.03 | 0.3905 |
| SPH | 1.94 ± 0.81 | 1.44 ± 0.31 | 0.1024 |
| S1P | 0.29 ± 0.23 | 0.44 ± 0.15 | 0.0481* |

**Supplementary Table 6.** Targeted sphingolipid abundance in the myometrium at term non-labor and term labor. Statistical significance was determined by two-way ANOVA, and significant differences were followed up with Holm-Šídák's multiple comparisons test. Data represented are mean ± SD. TNL: term non-labor; TL: term labor; SM: sphingomyelin; Cer: ceramide; SPA: sphinganine; SPH: sphingosine; S1P: sphingosine-1-phosphate.

**Supplemental Table 7**. Sphinganine, sphingosine, and S1P concentrations in the human chorioamnion, decidua parietalis, and myometrium at preterm non-labor and term non-labor.

| Sphingolipid (nmol/g tissue) | **PTNL (n=6)** | | | **TNL (n=8)** | | | **Adjusted p-Value PTNLvTNL** | | |
|---|---|---|---|---|---|---|---|---|---|
| | A | D | M | A | D | M | A | D | M |
| SPA[a,c,d,f] | 0.17 ± 0.12 | 0.68 ± 0.25 | 0.15 ± 0.06 | 0.24 ± 0.16 | 0.67 ± 0.30 | 0.28 ± 0.17 | 0.4239 | 0.9392 | 0.0745 |
| SPH[a,b,d,e] | 0.58 ± 0.30 | 2.00 ± 0.87 | 1.4 ± 0.70 | 0.65 ± 0.33 | 1.76 ± 0.49 | 1.94 ± 0.81 | 0.6765 | 0.5667 | 0.2110 |
| S1P[a,e,f] | 0.04 ± 0.01 | 0.1 ± 0.08 | 0.07 ± 0.03 | 0.08 ± 0.05 | 0.13 ± 0.08 | 0.29 ± 0.23 | 0.0467* | 0.4390 | 0.0301* |
| S1P:SPA[c,e,f] | 0.35 ± 0.35 | 0.15 ± 0.11 | 0.63 ± 0.51 | 0.6 ± 0.64 | 0.21 ± 0.1 | 1.2 ± 0.64 | 0.3754 | 0.3018 | 0.0847 |
| S1P:SPH | 0.08 ± 0.05 | 0.05 ± 0.02 | 0.06 ± 0.05 | 0.17 ± 0.16 | 0.07 ± 0.03 | 0.15 ± 0.09 | 0.1699 | 0.0927 | 0.0345* |

**Supplemental Table 7**. Significance → **a**: PTNL AvD; **b**: PTNL AvM; **c**: PTNL DvM; **d**: TNL AvD; **e**: TNL AvM; **f**: TNL DvM. Statistical significance was determined by two-way ANOVA, and significant differences were followed up with Holm-Šídák's multiple comparisons test. Data represented are mean ± SD. A: chorioamnion; D: decidua parietalis; M: myometrium; PTNL: preterm non-labor; TNL: term non-labor; SPA: sphinganine; SPH: sphingosine; S1P: sphingosine-1-phosphate.
