## Supplementary figures and images for "The Sphingosine-1-phosphate pathway is differentially activated in human gestational tissues"

### Expression of IL8 was used as a marker of labor and notably increased in TL myometrial samples (Supplementary Fig. 1)

A

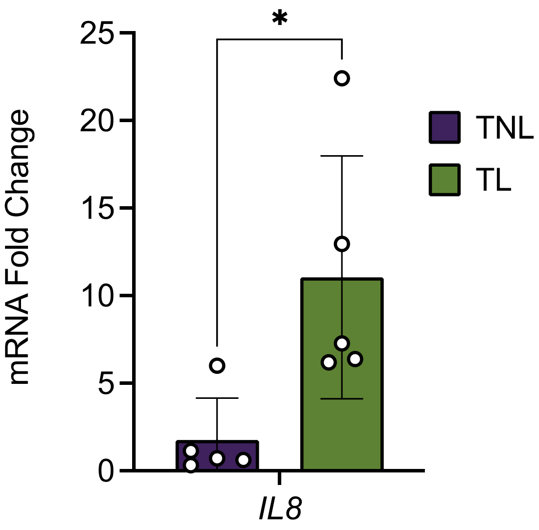
